# Refining epileptogenic high-frequency oscillations using deep learning: a reverse engineering approach

**DOI:** 10.1101/2021.08.31.458385

**Authors:** Yipeng Zhang, Qiujing Lu, Tonmoy Monsoor, Shaun A. Hussain, Joe X Qiao, Noriko Salamon, Aria Fallah, Myung Shin Sim, Eishi Asano, Raman Sankar, Richard J. Staba, Jerome Engel, William Speier, Vwani Roychowdhury, Hiroki Nariai

## Abstract

Intracranially-recorded interictal high-frequency oscillations (HFOs) have been proposed as a promising spatial biomarker of the epileptogenic zone. However, visual verification of HFOs is time-consuming and exhibits poor inter-rater reliability. Furthermore, no method is currently available to distinguish HFOs generated from the epileptogenic zone (epileptogenic HFOs: eHFOs) from those generated from other areas (non-epileptogenic HFOs: non-eHFOs). To address these issues, we constructed a deep learning (DL)-based algorithm using HFO events from chronic intracranial electroencephalogram (iEEG) data via subdural grids from 19 children with medication-resistant neocortical epilepsy to: 1) replicate human expert annotation of artifacts and HFOs with or without spikes, and 2) discover eHFOs by designing a novel weakly supervised model (HFOs from the resected brain regions are initially labeled as eHFOs, and those from the preserved brain regions as non-eHFOs). The “purification power” of DL is then used to automatically relabel the HFOs to distill eHFOs. Using 12,958 annotated HFO events from 19 patients, the model achieved 96.3% accuracy on artifact detection (F1 score = 96.8%) and 86.5% accuracy on classifying HFOs with or without spikes (F1 score = 80.8%) using patient-wise cross-validation. Based on the DL-based algorithm trained from 84,602 HFO events from nine patients who achieved seizure-freedom after resection, the majority of such DL-discovered eHFOs were found to be HFOs with spikes (78.6%, p < 0.001). While the resection ratio of detected HFOs (number of resected HFOs/number of detected HFOs) did not correlate significantly with post-operative seizure freedom (the area under the curve [AUC]=0.76, p=0.06), the resection ratio of eHFOs positively correlated with post-operative seizure freedom (AUC=0.87, p=0.01). We discovered that the eHFOs had a higher signal intensity associated with ripple (80-250 Hz) and fast ripple (250-500 Hz) bands at the HFO onset and with a lower frequency band throughout the event time window (the inverted T-shaped), compared to non-eHFOs. We then designed perturbations on the input of the trained model for non-eHFOs to determine the model’s decision-making logic. The model probability significantly increased towards eHFOs by the artificial introduction of signals in the inverted T-shaped frequency bands (mean probability increase: 0.285, p < 0.001), and by the artificial insertion of spike-like signals into the time domain (mean probability increase: 0.452, p < 0.001). With this DL-based framework, we reliably replicated HFO classification tasks by human experts. Using a reverse engineering technique, we distinguished eHFOs from others and identified salient features of eHFOs that aligned with current knowledge.

**Graphical abstract:** 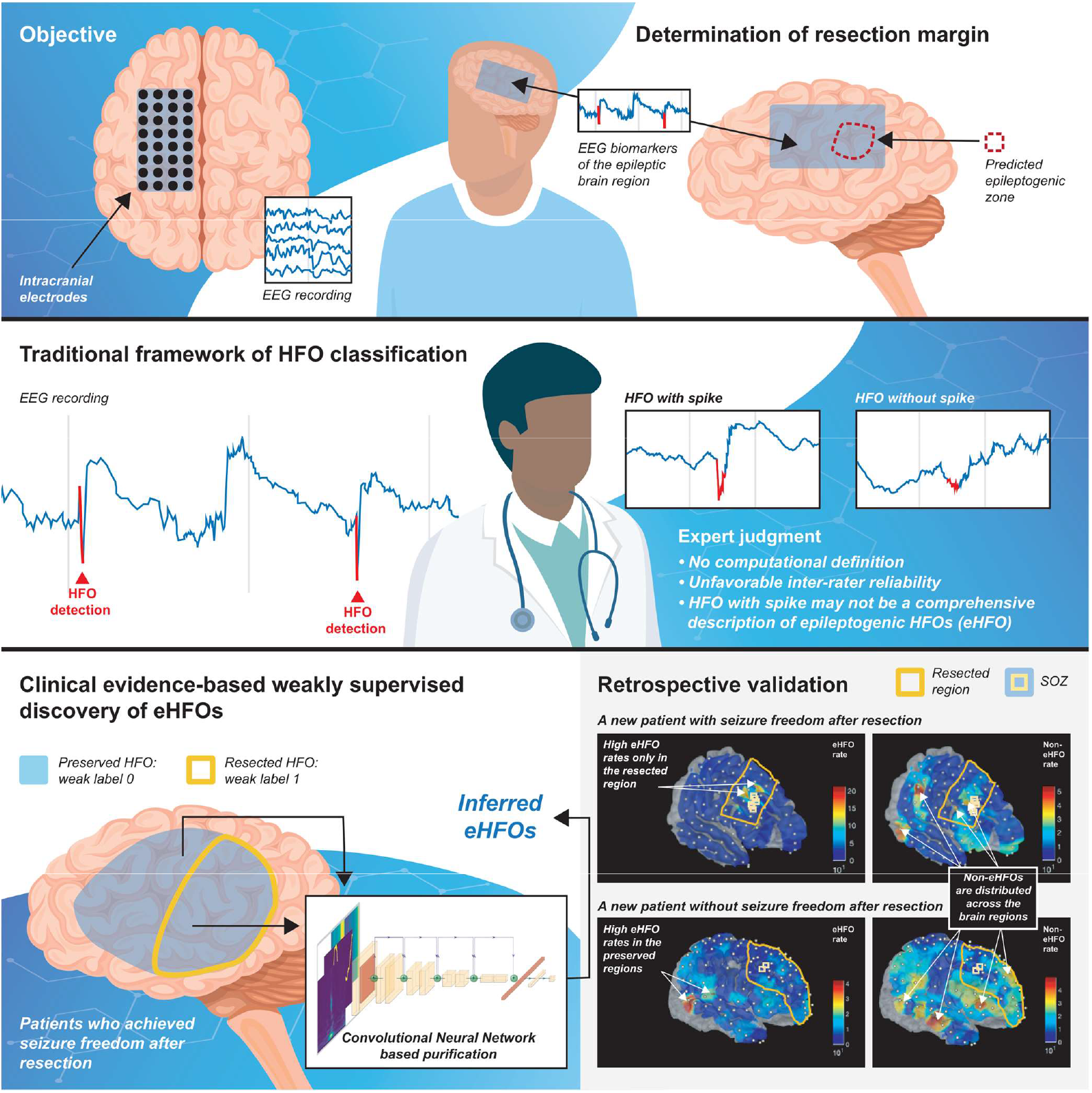

## INTRODUCTION

More than one-third of individuals with epilepsy are medication-resistant, making them potential surgical candidates.^1^ Currently, surgery is primarily guided by neuroimaging and neurophysiology (interictal spikes and seizure onset zone). However, the seizure-freedom rate of surgery is suboptimal, ranging from 50 to 85%.^2–5^ Identifying a biomarker that can accurately delineate the spatial extent of the epileptogenic zone (EZ: brain areas responsible for generating seizures) will be groundbreaking. Human and animal studies of epilepsy have suggested that intracranially-recorded interictal high-frequency oscillations (HFOs) on electroencephalogram (EEG) is a promising spatial neurophysiological biomarker of the epileptogenic zone.^6–10^ Many retrospective studies demonstrated that the removal of brain regions producing HFOs correlated with post-operative seizure-freedom.^11–14^ However, a recent prospective study could not reproduce these results.^15^ One of the most challenging issues is the presence of HFOs generated in healthy brain regions (physiological HFOs), which means seizure-freedom may be achieved despite leaving behind some areas displaying HFOs.^16–18^ In short, to utilize HFOs as a spatial biomarker to guide epilepsy surgery, one needs to establish a methodology to differentiate HFOs that are generated from the EZ (epileptogenic HFOs: eHFOs) and HFOs that are generated from other areas (non-epileptogenic HFOs: non-eHFOs).

A hypothesis-driven approach to look for eHFOs is quite challenging in practice because one needs to consider numerous and yet-to-be-identified features. Visual classification of HFOs with or without spikes along with artifact removal is commonly performed,^19^ because HFOs with spike-wave discharges are considered representative of eHFOs.^20–22^ However, this task is time-consuming and exhibits poor inter-rater reliability.^23^ Semiautomated computational methods to evaluate HFO characteristics (frequency, amplitude, and duration) do not appear to be useful for differentiating between eHFOs and non-eHFOs.^16, 24^ Fast ripples (250-500 Hz) might more specifically localize epileptogenic zones than ripples (80-250 Hz), but their detection rate is much lower than ripples.^12, 25, 26^ Correcting the HFO detection rate with region-specific normative values seems a reasonable approach,^27, 28^ but this does not determine each HFO event as an eHFO or non-eHFO.

Therefore, automated computational methods, such as those developed in the fields of artificial intelligence, would ideally discover eHFOs, guided purely by large samples of HFOs and clinical outcomes. Once trained, an ideal model should work robustly for any future patient. Moreover, these automated models should be interpretable to enable clinical decisions with high confidence and guide further scientific explorations of biological mechanisms. Indeed, machine learning has been successfully applied to the problem of classifying HFOs based on a priori manual engineering of event-wise features, which includes: linear discriminant analysis,^29^ support vector machines,^30, 31^ decision trees,^32^ and clustering.^33^ More recently, the deep learning (DL) framework has been adopted, which directly works with raw data (avoiding any a priori feature engineering) and yields better performance in the field of neuroimaging.^34^ Leveraging DL’s revolutionary success in the field of computer vision using Convolutional Neural Networks (CNNs), prior studies explored the use of CNNs in EEG analysis, especially converting one-dimensional EEG signal into a two-dimensional image for CNNs input.^35–37^ The previous DL approaches conducted the HFO classification in a supervised manner, requiring human annotated labels which constrains the spectrum of usage of their methods, especially the needs of human expert labeling. In the context of medical image analysis, recent work^38^ has shown that optimized model architectures and loss functions could mitigate data labeling errors, thus making the DL framework even more applicable.

The present study employed innovative analytic approaches to address several challenges expected in applying DL frameworks to the HFO classification task. As noted above, no direct observation of eHFOs is currently possible, making the most widely used supervised framework of DL impractical for our problem. Even if one were to solve this challenging problem, one still needs to make the DL models interpretable, which is yet another difficult task. In this study, we first proved that DL models could reliably emulate experts’ visual annotations in classifying the HFOs into artifacts, HFOs with spikes, or HFOs without spikes, without any a priori feature extractions. To mitigate potential labeling errors, we then generalized this approach to our central task of discovering eHFOs by replacing experts’ inputs with inexact weak labels implied by clinical outcomes and by using the “purification power” of DL to automatically distill eHFOs. Furthermore, (i) we proved the generalizability of this approach by using patient-wise cross-validation, implying a DL algorithm trained by EEG data from a large and diverse enough retrospective cohort is likely applicable to future patients; and (ii) we reverse engineered interpretable salient features of the DL-discovered eHFOs and showed that they aligned with current expert knowledge.

## METHODS

### Patient cohort

This was a retrospective cohort study. Children (below age 21) with medically refractory epilepsy (typically with monthly or greater seizure frequency and failure of more than three first-line anti-seizure medications) who had intracranial electrodes implanted for the planning of epilepsy surgery with anticipated cortical resection with the Pediatric Epilepsy Program at UCLA were consecutively recruited between August 2016 and August 2018. Diagnostic stereo-EEG evaluation (not intended for resective surgery) was excluded (**Table 1**).

**Table 1.** Cohort characteristics.

| Pt No. | Sex | Age at surgery (yr) | Epilepsy duration (yr) | Anti-seizure medications | No. of electrodes placed | No. of electrodes resected | % of electrodes resected | Duration of EEG (days) | No. of sz captured | MRI lesion/FDG-PET hypometabolism | Electrode coverage | Surgery | Pathology | No. of HFOs detected (10 min) | No. of HFOs detected (90 min) | Outcome (follow-up at 24 months) |
| --- | --- | --- | --- | --- | --- | --- | --- | --- | --- | --- | --- | --- | --- | --- | --- | --- |
| 1 | M | 20 | 6 | CLB, LVT, LCM | 40 | 9 | 22.50% | 3 | 21 | NL/ R FP | R FP | R focal resection of sensorimotor cortex | Gliosis | 334 | 1236 | Sz free |
| 2 | M | 11 | 9 | CLB, CNZ, LVT, RFD, PPN | 80 | 18 | 22.50% | 3 | 26 | R PO/ R PO | R FTPO | R focal resection around parietal tumor | Ganglioneurocytoma | 303 | 1673 | Sz free |
| 3 | F | 19 | 9 | LVT, LCM | 92 | 27 | 29.35% | 5 | 8 | R F/ R F | R FTP | R focal resection of frontal cortex | FCD 1b | 441 | 3433 | Sz free |
| 4 | F | 14 | 10 | CLB, LTG, LCM | 64 | 28 | 43.75% | 2 | 7 | L F/ L F | L FP | L frontal lobectomy sparing sensorimotor cortex | Gliosis | 593 | 5295 | Sz free |
| 5 | M | 9 | 7 | CLB, LTG | 100 | 83 | 83.00% | 6 | 4 | R TPO/ R TPO | R FTPO | R TPO | Gliosis | 513 | 2766 | Sz free |
| 6 | F | 3 | 2 | CLB, OXC | 96 | 42 | 43.75% | 2 | 18 | R F/ R FP | R FTP | R frontal lobectomy sparing sensorimotor cortex | FCD 2a | 280 | 2375 | Sz recurrence after 1 day |
| 7 | M | 5 | 3 | PB, PPN, OXC | 104 | 82 | 78.85% | 2 | 22 | L FP/ L FP | L FTP | L frontal lobectomy including resection of sensorimotor cortex | FCD 2a | 1177 | 9016 | Sz free |
| 8 | F | 19 | 7 | LVT, LCM | 104 | 0 | 0.00% | 2 | 23 | R TPO/ R TPO | R FTPO | No resection (RNS placement of R frontoparietal area) | NA | 481 | NA | NA |
| 9 | M | 13 | 6 | OXC, LTG, CLB | 72 | 5 | 6.94% | 6 | 7 | R P/ R P | R FP | R focal resection of parietal cortex | FCD 2a | 235 | 1384 | Sz recurrence after 10 days |
| 10 | F | 9 | 7 | CLB, OXC | 108 | 43 | 39.81% | 8 | 35 | L F/ L FP | L FTP | L frontal lobectomy sparing sensorimotor cortex | FCD 1c | 2921 | 25891 | Sz free for 20 months, then lost follow-up |
| 11 | F | 8 | 7 | LVT, LCM, CLB | 66 | 14 | 21.21% | 6 | 1 | L T/ L TP | L FTP | L temporal lobectomy | Multinodular and vacuolating neuronal tumor (MVNT) | 250 | 1670 | Sz free |
| 12 | F | 18 | 17 | CLB, LTG | 84 | 12 | 14.29% | 3 | 25 | L P/ L P | L FTP | L focal resection of parietal cortex | Gliosis | 268 | 1943 | Sz recurrence after 3 days |
| 13 | F | 15 | 3 | LCM, LVT, OXC | 86 | 9 | 10.47% | 4 | 4 | R F/ R F | R FTP | R focal resection around frontal tumor | Oligodendroglioma | 164 | 2243 | Sz free |
| 14 | F | 19 | 18 | LTG, LCM, OXC, PPN | 70 | 0 | 0.00% | 4 | 37 | R FP/ R FP | R FTPO | No resection (RNS placement of R frontoparietal area) | NA | 722 | NA | NA |
| 15 | F | 15 | 15 | TPM, LTG | 102 | 60 | 58.82% | 4 | 4 | L TPO/ L TPO | L FTPO | L TPO | FCD 2a, Gliosis | 1211 | 6867 | Sz free |
| 16 | M | 6 | 6 | CLB, OXC | 104 | 41 | 39.42% | 11 | 2 | L PO/ L PO | L FTPO | L parietooccipital resection | Ulegyria, FCD 3d, gliosis | 1920 | 15782 | Sz recurrence after 23 days |
| 17 | M | 20 | 15 | LCM, BVC, FBM | 118 | 29 | 24.58% | 12 | 5 | L TO/ L TO | L FTPO | L temporal lobectomy plus RNS | Gliosis | 465 | 3028 | Sz recurrence after 4 days |
| 18 | M | 12 | 5 | CNZ, CLP, ECZ, LCM, LVT | 112 | 0 | 0.00% | 4 | 100 | L P/ L TP | L FTPO | No resection (RNS placement of L sensorimotor cortex) | NA | 220 | NA | NA |
| 19 | M | 14 | 6 | ECZ, CLB | 128 | 0 | 0.00% | 14 | 8 | L T/ L TP | LFTPO | No resection (RNS placement of L temporal, parietal, and occipital area) | NA | 469 | NA | NA |

M: Male; F: Female; FRs: Fast ripples; NA: Not applicable; RNS: Responsive nerve stimulator; FCD: Focal cortical dysplasia; SOZ: Seizure onset zone; Sz: Seizure.
L: Left; R: Right; F: Frontal; P: Parietal; T: Temporal; O: Occipital.
CLB: Clobazam; LVT: Levetiracetam; LCM: Lacosamide; CNZ: Clonazepam; RFD: Rufinamide; PPN: Perampanel; LTG: Lamotrigine; OXC: Oxcarbazepine; PB: Phenobarbital;
TPM: Topiramate; BVC: Brivaracetam; FBM: Felbamate; CLP: Clorazepate; ECZ: Eslicarbazepine.

### Standard protocol approvals, registrations, and patient consents

The institutional review board at UCLA approved the use of human subjects and waived the need for written informed consent. All testing was deemed clinically relevant for patient care, and also all the retrospective EEG data used for this study were de-identified before data extraction and analysis. This study was not a clinical trial, and it was not registered in any public registry.

### Patient evaluation

All children with medically refractory epilepsy referred during the study period underwent a standardized presurgical evaluation, which—at a minimum—consisted of inpatient video-EEG monitoring, high resolution (3.0 T) brain magnetic resonance imaging (MRI), and 18 fluoro-deoxyglucose positron emission tomography (FDG-PET), with MRI-PET co-registration.^26^ The margins and extent of resections were determined mainly based on seizure onset zone (SOZ), clinically defined as regions initially exhibiting sustained rhythmic waveforms at the onset of habitual seizures. In some cases, the seizure onset zones were incompletely resected to prevent an unacceptable neurological deficit.

### Subdural electrode placement

Macroelectrodes, including platinum grid electrodes (10 mm intercontact distance) and depth electrodes (platinum, 5 mm intercontact distance), were surgically implanted. The total number of electrode contacts in each subject ranged from 40 to 128 (median 96 contacts). The placement of intracranial electrodes was mainly guided by the results of scalp video-EEG recording and neuroimaging studies. All electrode plates were stitched to adjacent plates, the edge of the dura mater, or both, to minimize movement of subdural electrodes after placement.

### Acquisition of three-dimensional (3D) brain surface images

We obtained preoperative high-resolution 3D magnetization-prepared rapid acquisition with gradient echo (MPRAGE) T1-weighted image of the entire head. A FreeSurfer-based 3D surface image was created with the location of electrodes directly defined on the brain surface, using post-implant computed tomography (CT) images.^39^ In addition, intraoperative pictures were taken with a digital camera before dural closure to enhance spatial accuracy of electrode localization on the 3D brain surface. Upon re-exposure for resective surgery, we visually confirmed that the electrodes had not migrated compared to the digital photo obtained during the electrode implantation surgery.

### Intracranial EEG (iEEG) recording

Intracranial EEG (iEEG) recording was obtained using Nihon Kohden Systems (Neurofax 1100A, Irvine, California, USA). The study recording was acquired with a digital sampling frequency of 2,000 Hz, which defaults to a proprietary Nihon Kohden setting of a low frequency filter of 0.016 Hz and a high frequency filter of 600 Hz at the time of acquisition. For each subject, separate 10-minute and 90minute EEG segments from slow-wave sleep were selected at least two hours before or after seizures, before antiseizure medication tapering, and before cortical stimulation mapping, which typically occurred two days after the implant. All the study iEEG data were part of the clinical EEG recording.

### Automated detection of HFOs

A customized average reference was used for the HFO analysis, with the removal of electrodes containing significant artifacts. Candidate interictal high frequency oscillations (HFOs) were identified by an automated short-term energy detector (STE).^40, 41^ This detector considers HFOs as oscillatory events with at least six peaks and a center frequency occurring between 80-500 Hz. The root mean square (RMS) threshold was set at five standard deviations (SD), and the peak threshold was set at three SD. The HFO events are segments of EEG signals with durations ranging from 60 to 200 ms (see SI for duration distribution). We referred to these detected events as candidate HFOs (c-HFOs).

### Human expert classification of HFOs

A human expert (HN: board certified in clinical neurophysiology and epilepsy, with experience in HFO analysis) classified c-HFOs in each patient’s 10-minute EEG segments into three classes: HFOs with spikes (spk-HFO), HFOs without spikes (non-spk-HFO), and artifacts using RippleLabs graphic user interface,^41^ based on three images (unfiltered EEG tracing, filtered EEG tracing [80-500 Hz], and time-frequency plot). Artifacts are false-positive events, including ringing (filtering of sharp transients),^19^ as well as muscle and background fluctuations (see examples in **Figure 1**). Another expert with similar qualifications (SH) independently scored c-HFOs from two representative patients, and inter-rater reliability was examined using Cohen’s kappa statistics.

**Figure 1.**
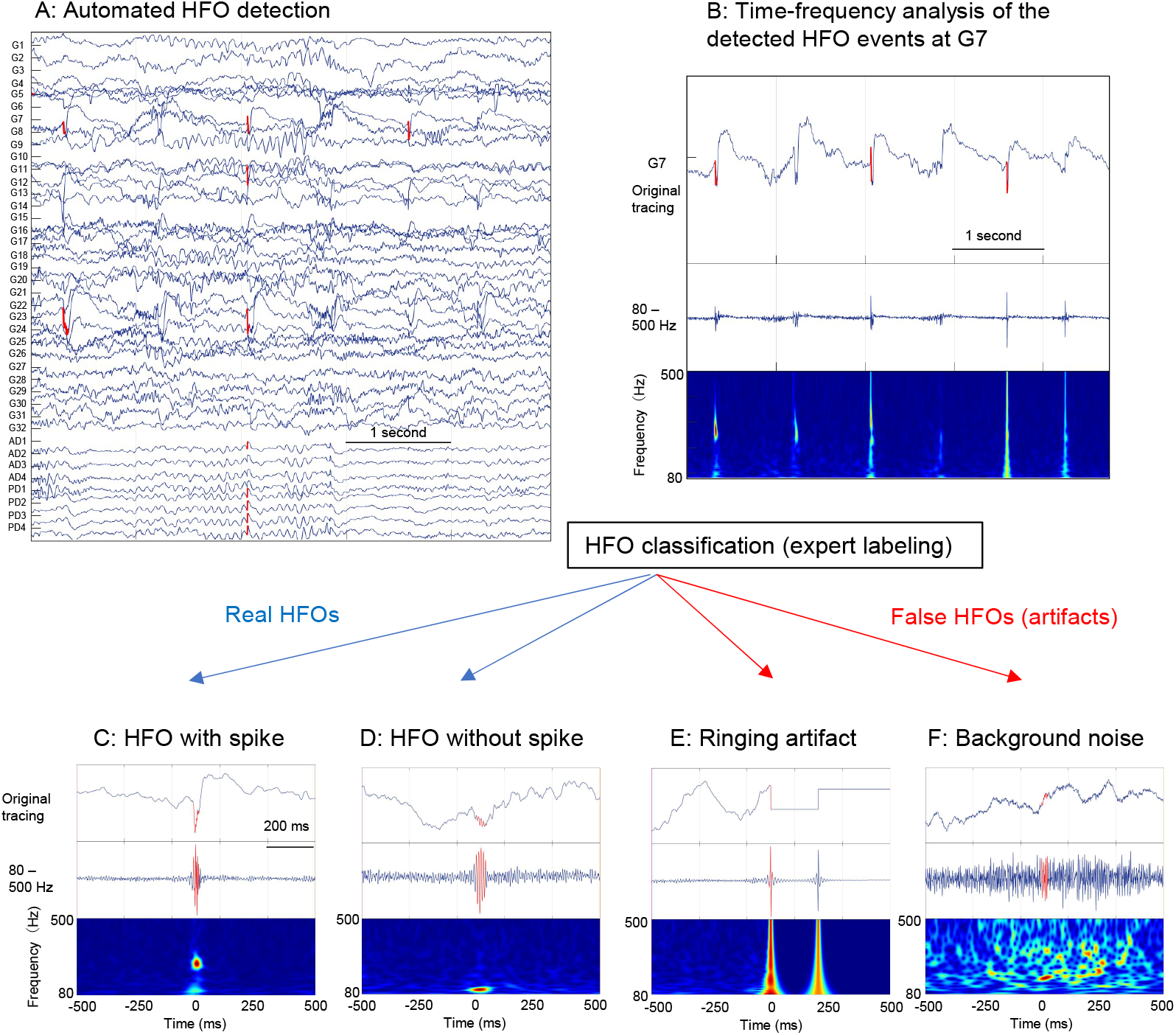
Automated detection of HFOs and classification of HFOs by a human expert. After each EEG sample was arranged with a referential montage, the short-term energy (STE) HFO detector was applied to detect candidate HFO events (A). The detected HFO events were marked in the original tracing (B). Each detected HFO event was reviewed by a human expert to classify into HFO with spike (spk-HFO) (C), HFO without spike (D), and artifacts: ringing artifact (E) and background noise (F). EEG: Electroencephalogram; HFOs: High-frequency oscillations; STE: Short-term energy

### Supervised deep learning networks using expert labels

The general workflow of the DL training and inference were shown in the flowchart (**Figure 2**)

**Figure 2.**
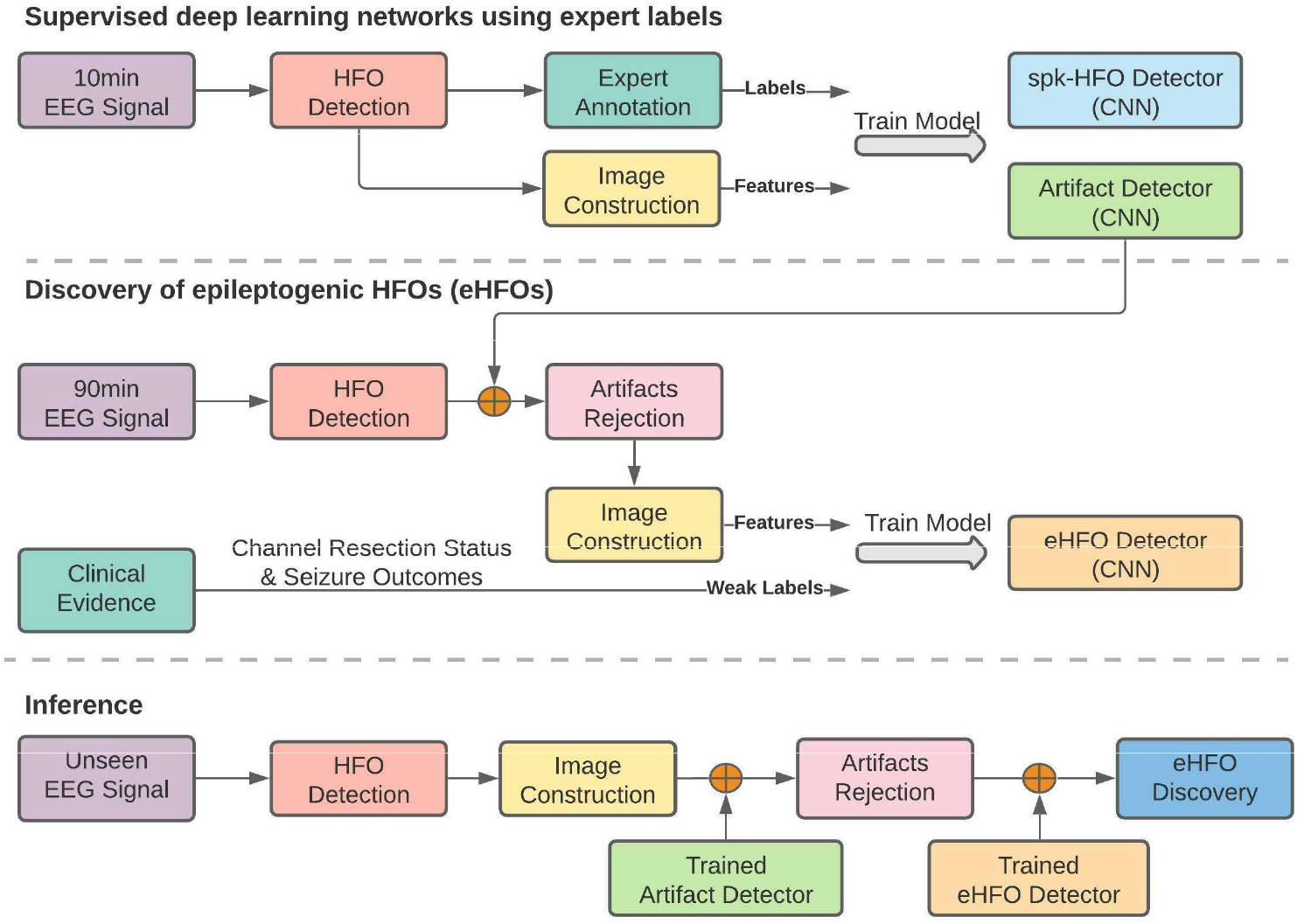
Processing workflow. Our study’s overall data processing workflow is shown as a flowchart.

#### Feature representation of c-HFOs

Each c-HFO was represented by a one-second window, with the c-HFO was located at the center (0 ms), and including 500ms of EEG signal before and after. To utilize the power of CNN, we captured the time-frequency domain features as well as signal morphology information of the c-HFO window via three images (**Figure 3A**). The <u>time-frequency plot</u> (scalogram) was generated by continuous Gabor Wavelets ranging from 10Hz to 500Hz.^41^ The <u>EEG tracing plot</u> was generated on a 2000 x 2000 image by scaling the timeseries signal into the 0 to 2000 range to represent the EEG waveform’s morphology. The <u>amplitude-coding plot</u> was generated to represent the relative amplitude of the time-series signal: for every time point, the pixel intensity of a column of the image represented the signal’s raw value at that time. These three images were resized into the standard size (224 x 224), serving as the input to the neural network.

**Figure 3:**
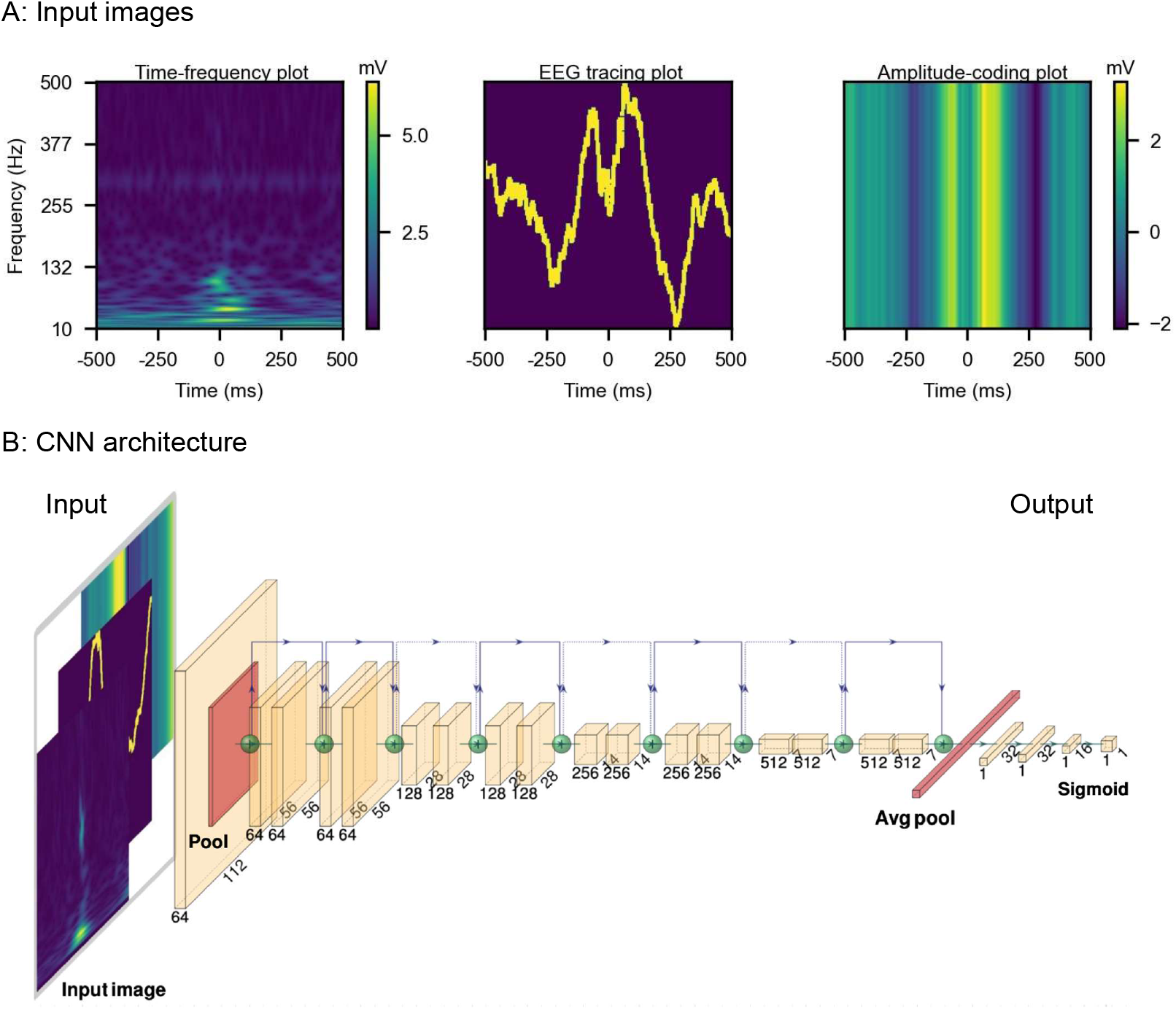
Network input and architecture. (A) Network input images. The network input includes three images constructed from a one-second raw EEG segment with a detected HFO in the center (500ms before and after). Left: The time-frequency plot was generated by continuous Gabor Wavelets ranging from 10Hz to 500Hz. Middle: EEG tracing plot was generated on a 2000 x 2000 image by scaling the time-series signal into the 0 to 2000 range. Right: amplitude-coding plot contains the amplitude at every time point; a column of the image represented the signal’s actual value rescaled with a color gradient. These three images were resized into 224 x 224 in order to fit into the neural network. (B) CNN architecture. The architecture of the model was adapted from Resnet-18. The last layer of the resnet18 was modified to be three fully-connected layers with LeakyReLU, BatchNorm, and 10 % dropout in between. The output of the model was fed into a sigmoid function to bound the output between 0 to 1, representing the probability of each task. For task 2 (HFOs with spikes vs. HFOs without spikes), the input consisted of the three images (time-frequency plot, EEG tracing plot, and amplitude-coding plot). Meanwhile, for task 1 (Real HFOs vs. artifacts), only the time-frequency plot information was found to be sufficient, and hence three same time-frequency plots were concatenated together as the input of the model.

#### Two-step deep learning model architecture

Given the expert labeling with three labels (artifacts, spk-HFO, and non-spk-HFO), the training of the deep neural network (DNN) model was formulated as two binary classification steps. Step1 (artifact detector): we differentiated between artifacts and “Real HFOs”, defined as the union of spk-HFOs and non-spk-HFOs. All “Real HFOs” were labeled as the positive samples and the artifacts were the negative samples. Step2 (spk-HFO detector) classified the “Real HFOs” into spk-HFO and non-spk-HFO; the spk-HFO were defined as positive samples, and the non-spk-HFO were defined as negative samples.

The artifact detector and spike detector’s architectures are identical and adapted from ResNet18^42^ with a modification in the last few layers to accommodate the binary classification tasks. Specifically, the last layer of the resnet18 was modified to be three fully-connected layers with LeakyReLU, BatchNorm, and 10 % dropout in between. The output of the model was fed into a sigmoid function to bound the output between 0 to 1, representing the probability of each task. The input comprising three image channels and the architecture of the networks are shown in **Figure 3B**. For the artifact detector, only the time-frequency information was used. Hence time-frequency plots were repeated three times and concatenated together as the input to the artifact detector. For the spk-HFO detector, concatenation of the three feature-representing images (time-frequency plot, EEG tracing plot, and amplitude-coding plot) served as input.

#### Training and performance analysis

There were two types of training conducted: patient-wise cross-validation and all-patients training. For patient-wise cross-validation, one patient was selected at a time as the test set, and the remaining patients were used for model training. All events were pooled across the rest of the patients, with 10% randomly sampled to serve as a validation set and the remaining 90% used for training. In all-patients training, five-fold cross-validation was conducted across the pooled data across the full patient cohort. For each fold, 20% of the dataset was selected as the test set, 70% was selected as the training set, and the remaining 10% was used for validation.

Since the optimization goal of both detectors is binary classification, we adopted binary cross-entropy as the loss function and the Adam optimizer^32^ with a learning rate of 0.0003. All of the training was conducted using 15 epochs (training iterations) and validation loss was plotted with respect to the number of epochs completed. For the artifact detector, to improve generalization, we picked the model in the epoch that corresponds to the first local minima in the plot; this technique is also known as an early stopping regularization. For the spike detector, we directly picked the model corresponding to the global minima over 15 iterations, i.e., the lowest validation loss.

We calculated the precision, recall, and accuracy of the classification results. To measure the model performance on an unbalanced dataset like ours, we also calculated the F1-score. For the patient-wise cross-validation task, we averaged the performance statistics across patients. For the all-patients training task with five-fold crossvalidation, we reported average model performance over cross-validation folds.

### Discovery of epileptogenic HFOs via deep learning based on clinical outcomes and channel resection status

While no direct observation of eHFOs is currently possible, clinical evidence such as seizure outcomes and resection status of the channels can be used to determine highly likely groups of eHFOs and non-eHFOs. Such data-driven inexact or weak labels thus contain unknown labeling errors and in order to further purify the labels, one would need to do an automated geometric similarity analysis in the HFO feature space: the HFOs that are geometric outliers in the group dominated by eHFOs should be relabeled as non-eHFOs and vice versa. A universal classifier, such as a DNN, has the potential to do exactly this when trained with weak labels: it automatically computes an optimal boundary in the space of all HFOs so that the geometric outliers get separated by this boundary. The trained DNN classifier then could determine when any given HFO is most likely an eHFO and its confidence level. This blackbox classifier was then probed and analyzed to obtain a more computational definition of eHFOs, a step akin to reverse engineering.

#### Label assignment for training: Weak Supervision

We generated a weakly labeled training set with the following assumptions. For those patients that became seizure-free after resection (9 patients), we assumed that all epileptogenic tissue was contained in the resection. Similarly, the preserved regions in post-surgery seizure-free patients could not contain epileptogenic tissue, leading us to assume that the HFOs in the preserved regions are non-epileptogenic. We thus labeled all of the HFOs from resected channels as 1 and all HFOs from preserved channels as 0. By doing such weak supervision, we introduced potential errors in the data labels. Specifically, (1) the epileptogenic zone may generate non-eHFOs in addition to eHFOs; (2) some tissue was resected based on anatomical location (e.g. the channel was located within the same gyrus as the SOZ) and was likely not epileptogenic, leading to false-positive labels; and (3) in the non-resected region, while the majority of the HFOs were indeed non-epileptogenic, a few HFOs that are morphologically similar to epileptogenic HFOs could have been introduced by phenomena such as propagation.^43^ To address (1) and (3), we believed the automatic geometric purification property of the neural network could denoise the errors introduced by this mislabeling since the clinical outcomes constrain the number of mislabels to be few. To address (2), we introduced a weight term in the network’s loss function to reduce the noise introduced by the false positives, which is described in the following section. Note that for patients that were not seizure-free after surgery, the assumption that all epileptogenic tissue lay within the resected boundary did not hold. Thus, they were not used for any training.

To maximize the training set size, we used 90 minutes of data from each patient for training. In order to ensure that our proposed framework generalizes across patients, a patient-specific model was designed for each patient without using any of its data. Thus, for a post-operative seizure-free patient, a patient-specific model was trained on data from other patients who became seizure-free after surgery. For post-operative non-seizure-free patients, a patient-specific model was trained on data from all post-operative seizure-free patients. Specifically, for training, we used 90 minutes of EEG data from 8 patients (if the target patient is post-surgery seizure-free) or 9 patients (if the target patient is not post-surgery seizure-free). All predictions (inference) used for the rest of the analysis were generated following this strategy.

#### The Deep Learning Architecture and Training for REVerse engineering (DLATREV)

The training details are as follows. We first used the artifact detector trained on all patients’ 10 min EEG data to filter out the potential artifacts in the c-HFOs from the 90 min EEG data. For the sake of convenience, any HFOs that passed this filtering, and hence are “Real HFOs,” would simply be referred to as HFOs going forward. The total number of HFOs differed considerably among the patients, and a data balancing process was required to balance the information introduced from each patient. For each patient, if the number of HFOs was smaller than a threshold of 2,500 (the median of the HFO distribution among patients), all of the HFOs were used. Otherwise, 2,500 events were uniformly sampled from that patient without replacement. There could potentially be many non-eHFOs in the non-seizure-onset but resected channels that were labeled incorrectly as eHFOs in our weak supervision process. For each input with a label of y and prediction of x, we introduced a weight term, **w**, in the Binary Cross Entropy (BCE) loss. (Loss = w* BCE Loss, where BCE Loss = −[y·log(x)+(1−y)·log(1−x)]). If y = 0 (the HFO is from a preserved channel), then w = 1. If y = 1 (the HFO is from a resected channel) and the HFO is from a SOZ channel, then also w = 1. However, if y = 1 and the HFO is not from a SOZ channel, we used a value of w = 0.5, which was found to be optimal for our dataset (See **Supplementary Figure**). For the model architecture and training, we used the same setup as the spike detector, except the CNN’s convolution layers were frozen to only act as feature extractors.

#### Relationship between epileptogenic HFOs and HFOs with spikes

In order to understand the correspondence between the discovered eHFOs or non-eHFOs and common clinical knowledge about such HFOs (i.e. HFOs with and without spikes), we let the model classify each HFO event on the 10 minutes annotated data using DLATREV. We quantitatively analyzed the correlation of classified eHFOs and non-eHFOs with their corresponding spk-HFO annotation using the chi-square test.

#### Spatial distribution of epileptogenic and non-epileptogenic HFOs

To visualize the distribution of the eHFOs and non-eHFOs in each patient’s data, we performed inference on HFOs obtained from 90-minute EEG data from 15 patients who underwent resection, using their corresponding patient-specific DLATREV models. Then we plotted the voltage map on the 3D cortical reconstruction to visualize the performance based on the number of eHFOs and non-eHFOs using the FreeSurfer-based cortical modeling.^22, 39^ The rate of eHFOs and non-eHFOs was compared in the SOZ and non-SOZ.

#### Comparison of resection ratios of HFOs to post-operative seizure outcomes

We estimated the probability of each patient’s surgical success (for the 14 patients who underwent resective surgery with known seizure outcome at 24 months) based on the resection ratio of HFOs (number of resected HFOs/ number of detected HFOs) as a classifier. We constructed the receiver operating characteristic (ROC) curve and calculated the area under the curve (AUC) values in the resection ratio of 1) c-HFOs, 2) Real HFOs, and 3) eHFOs. Determination of the channel resection status (resected vs. preserved) was determined based on intraoperative pictures (pre-and postresection) and also on post-resection brain MRI, based on discussion among a clinical neurophysiologist (HN), neurosurgeon (AF), and radiologist (SN). A multiple logistic regression model incorporating the resection ratio of eHFOs and complete resection of the SOZ was also created. The surgical outcomes were determined 24 months after resection, as either seizure-free or not seizure-free.

#### Time-frequency plot characteristics of epileptogenic and non-epileptogenic HFOs

We determined whether the time-frequency scalogram of eHFOs differed from that of non-eHFOs. This comparison constituted the first step in reverse engineering the computations that the DLATREV model learned to perform while executing its classification task. For every pixel (x,y) in a 224*224 image, we created two sets of data points, SeHFO (x,y) and Snon-eHFOs (x, y). SeHFOs(x,y) consisted of the intensity values f(x,y) of the scalogram for all the classified eHFOs for all patients. Similarly, Snon-eHFOs (x, y) consisted of the intensity values f(x,y) of the scalogram for all the classified non-eHFOs for all patients. Then we performed one-tailed t-tests to determine whether a random variable A(x,y), whose samples are given by SeHFO (x,y), is greater than that of a random variable B(x,y) whose samples are given by Snon-eHFOs (x, y). If this hypothesis was returned to be true with a p-value less than 0.005, we set the pixel value I(x,y) = 1 otherwise I(x,y) = 0.

#### Perturbation analysis to investigate salient features of epileptogenic and non-epileptogenic HFOs

There exist several potential ways to determine an interpretable function computed by a blackbox classifier such as a DNN, including (i) training an interpretable model with the outputs of the classifier,^44^ and (ii) adversarial perturbations of the inputs so as to effect maximum changes in the probability of the predicted outcomes such as Grad-CAM.^45^ Both of these methods are computationally expensive, and may not yield useful answers without prior knowledge. The knowledge distilled from the time-frequency plot characteristics of eHFOs and non-eHFOs and positive correlation between the eHFOs and spk-HFO was used for the perturbation analysis. We hence conducted the following two perturbations on all classified non-eHFOs in 90 min data to investigate the salient features that are critical to the model’s decision.

##### Perturbation on time-frequency plot

The I(x,y) image provided a template of the pixels where eHFOs had statistically significantly higher magnitudes than non-eHFOs. Then we designed an inverted T-shaped mask to approximate this template. We hypothesized that if we perturbed the scalogram of a non-eHFO over the template using the maximum value of the corresponding scalogram (thus making the scalogram similar to that of an eHFO), then the probability of the classifier output should significantly increase, making it look more like an eHFO to the classifier. In practice, each new value within this template equals 0.5 * original value + 0.5 * maximum value. Note that this method perturbed only the scalogram channel input and kept the other channel inputs unchanged. We summarized the results via a histogram of the change in probabilities for each patient across all classified non-eHFOs. Additionally, a one-tailed t-test comparing values of eHFOs and non-eHFOs, was performed on the change of output probability score to ensure that the change was significant and generalized well on the population level.

##### Perturbation on amplitude-coding plot

We hypothesized that a spike-like pattern close to an HFO detection in the time domain was a salient feature contributing to the eHFO discovery. We referred to this pattern as the upgoing or downgoing time-domain characteristic pattern (TDCP). There were two channels in which we could inject a TDCP: the EEG tracing plot and the amplitude-coding plot. We picked the amplitude-coding plot as it contained denser information than the EEG tracing plot, so that the model can have a more evident response to the perturbation. We mimicked the insertion of a TDCP by centering it at a particular time step and then replacing the values in the corresponding columns of the amplitude-coding with that of the TDCP. Furthermore, we scaled the TDCP such that for an upgoing TDCP, its peak equals the maximum value in the whole plot, and similarly for the downgoing TDCP, its valley equals the minimum value in the whole plot. In the perturbation, we kept the value of other channels the same. If the hypothesis were true, we would expect the perturbed amplitude-coding input for a non-eHFO to significantly increase the classifier output probability, making it look more like an eHFO to the classifier. We searched through all possible placements of the TDCP in the amplitude-coding plot to determine the location that led to the most perturbation to the model. The column (where the center of the TDCP was placed) with the maximum change of the probability score was viewed as the optimal time-location of the TDCP. We summarized the results via the histogram of the change in probabilities for each patient across all the classified non-eHFOs. A one-tailed t-test was also performed on the change of that output probability score to ensure that the change was significant and generalized well on the population level.

### Statistical analysis

Above mentioned statistical calculations were carried out using Python (version 3.7.3; Python Software Foundation, USA) and JMP Pro (version 14; SAS Institute, USA). The deep neural network was developed using PyTorch (version 1.6.0; Facebook’s AI Research lab). Quantitative measures are described by medians with interquartile, or means with standard deviations. Comparisons between groups were performed using chi-square for comparing two distributions and Student’s t-test for quantitative measures (in means with standard deviations). All comparisons were two-sided and significant results were considered at p < 0.05 unless stated otherwise. Specific statistical tests performed for each experiment were described in each section. Machine learning model performance was evaluated using accuracy ([TP + TN]/[TP+TN+FP+FN]), recall (TP/[TP+FN]), precision (TP/[TP+FP]), and F-1 score (2/[1/recall + 1/precision]).

### Data sharing and availability of the methods

Anonymized EEG data used in this study are available upon reasonable request to the corresponding author. The python-based code used in this study is freely available at (https://github.com/roychowdhuryresearch/HFO-Classification). One can train and test the deep learning algorithm from their data and confirm our methods’ validity and utility.

## RESULTS

### Patient characteristics

There were 19 patients (10 females) enrolled during the study period. The median age at surgery was 14 years (range: 3-20 years). Median electrocorticography monitoring duration was 4 days (range: 2-14 days), and the median number of seizures captured during the monitoring was 8 (IQ range: 4-25). There were 15 patients who underwent resection, and 14 patients provided post-operative seizure outcomes at 24 months (9 of 14 became seizure-free). Details of patients’ clinical information are listed in **Table 1**.

### Interictal HFO detection

Two experts showed favorable inter-rater reliability when 583 c-HFOs were labeled independently (kappa = 0.96 for labeling artifacts, 0.85 for labeling HFOs with spikes); thus, labels from one expert (HN) were used for the rest of the study. A total of 12,958 HFO events were detected (median 456 events per patient) in 10-minute EEG data from the 19 patients. The expert classification yielded 6,430 HFOs with spikes, 3,721 HFOs without spikes, and 2,807 artifacts. The 90-minute EEG data from the nine patients who became seizure-free after 24 months yielded 34,199 HFO events in total (median 2,570.5 events per patient). The 15 patients who had 90-minute data yielded 84,602 HFO events in total (median 2,766 events per patient).

### Machine learning algorithm against expert labeling

In the patient-wise cross-validation, our model achieved 96.3% accuracy [std = 4.96] on artifact detection (recall = 98.0% [std = 2.16%], precision = 96.1% [std = 8.72%], F1 score = 96.8% [std = 5.12%]) and 86.5% accuracy [std = 8.33%] for detecting HFOs with spikes (recall = 83.7% [std = 17.06%], precision = 81.4% [std = 16.07%], F1 score = 80.8% [std = 16.63%]). In all-patients 5-folds training, the model achieved accuracy of 98.9% [std = 0.15%] and 89.1% [std = 1.00%] for detecting artifacts and HFO with spikes, respectively. Additionally, our trained models achieved F1 score of 99.3% [std = 0.33%] (recall = 99.2% [std = 0.40%], precision = 99.4% [std = 0.40%]) and F1 score of 89.1% [std = 2.12%] (recall = 91.0% [std = 0.89%], precision = 92.0% [std = 0.84%]) for the two classification tasks.

### Relationship between eHFOs and HFOs with spikes

The DLATREV model (trained on 90-minute data) was applied to the 10-minute EEG dataset to discover eHFOs. These results were compared with the spk-HFO labels annotated by an expert. We noted 71.1% (4573/6430) of the eHFOs were HFOs with spikes, and 73.7% (2739/3721) of non-eHFOs were HFOs without spikes (p < 0.0001, a chi-square test).

### Spatial mapping of eHFOs and non-eHFOs

The DLATREV model was applied on a 90-minute EEG dataset for all subjects who underwent resection (n = 15) and the classified eHFOs and non-eHFOs were mapped on each reconstructed 3D MRI (examples in **Figure 4A**). In general, eHFOs clustered around the SOZ, while non-eHFOs distributed diffusely. The rate of eHFOs (eHFOs/min/channel) was higher in SOZ than that in non-SOZ (mean 0.74 vs. 0.28, p = 0.02, paired t-test), and the rate of non-eHFOs (non-eHFOs/min/channel) did not differ between the SOZ and non-SOZ (mean 0.36 vs. 0.29, p = 0.27, paired t-test) (**Figure 4B**).

**Figure 4.**
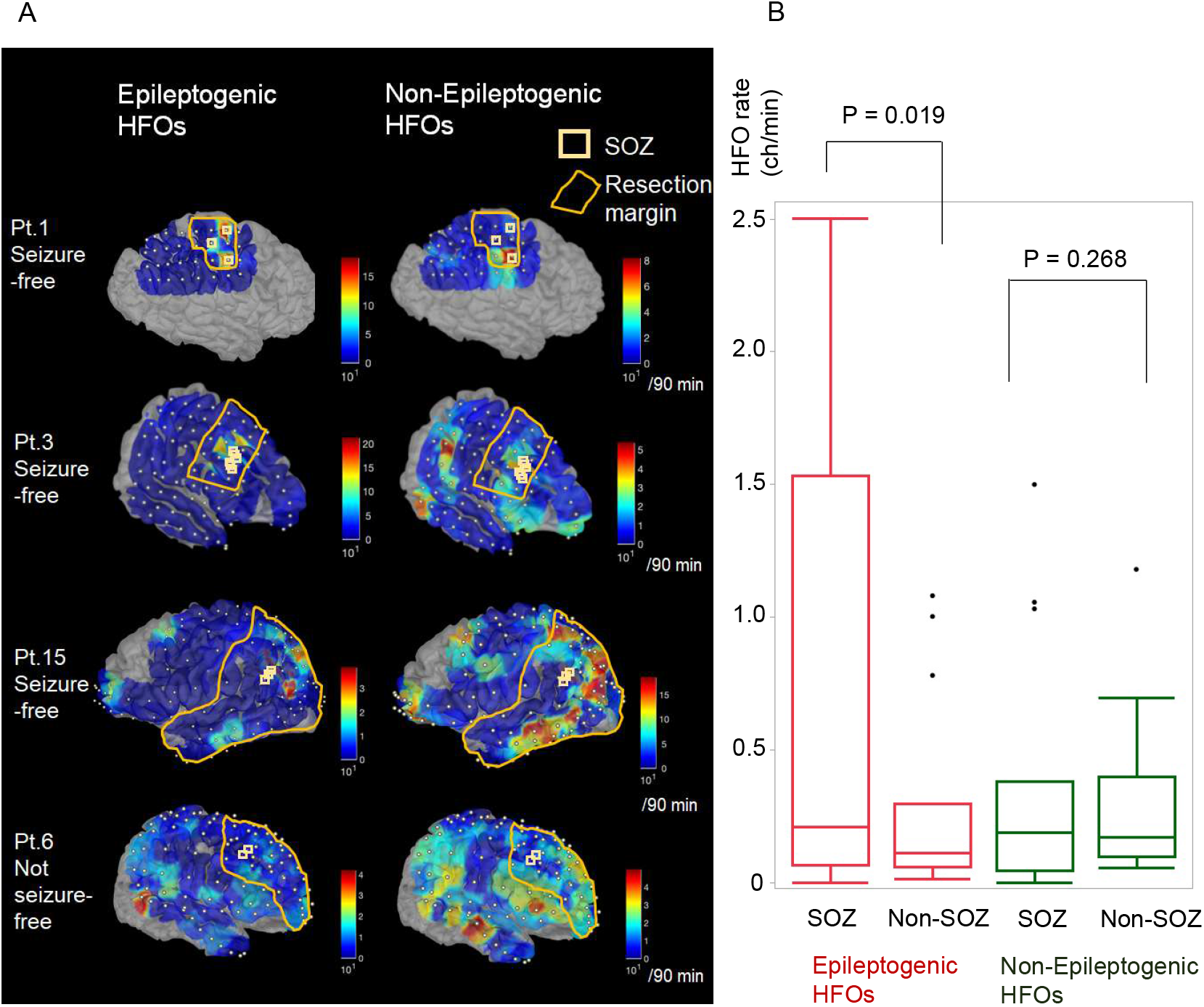
Spatial mapping of epileptogenic and non-epileptogenic HFOs by the DLATREV model. (A) In each row, individual 3D-reconstructed MRI with spatially coregistered electrodes is shown. Using a model developed using patient-wise cross-validation, the number of epileptogenic and non-epileptogenic HFOs (number/90 minutes) are projected onto the individual 3D MRI as an inference. Seizure onset zones are marked with white squares, and resected brain regions are marked with orange lines. The first three rows include subjects who achieved seizure-freedom after resection. The last row represents a subject who did not achieve seizure-freedom after resection. Epileptogenic HFOs localize around the seizure onset zones, whereas non-epileptogenic HFOs are localized more diffusely in the entire hemispheres. (B) The rate of HFOs (epileptogenic and non-epileptoenic HFOs) in each patient (n=15) is plotted in box plots based on the location (SOZ vs. non-SOZ). The rate of eHFOs (eHFOs/min/channel) was higher in SOZ than that in non-SOZ (mean 0.74 vs. 0.28, p = 0.02, paired t-test). The rate of non-eHFOs (non-eHFOs/min/channel) did not differ between the SOZ and non-SOZ (mean 0.36 vs. 0.29, p = 0.27, paired t-test) HFOs: High-frequency oscillations; SOZ: Seizure onset zone; Pt: Patient; MRI: Magnetic resonance imaging.

### Prediction of post-operative seizure-outcomes using the HFO classification algorithms

We created the ROC curves using HFO resection ratio to predict post-operative seizure freedom at 24 months (n = 14) (**Figure 5**). Using the resection ratio of c-HFOs and Real HFOs showed acceptable prediction performance but did not show statistical significance (AUC=0.78 and 0.76; p = 0.05 and 0.06, respectively). The use of resection ratio of eHFOs exhibited a high AUC value of 0.87 (p = 0.01). The performance was further augmented by using a multiple regression model incorporating both the resection ratio of eHFOs and complete removal of SOZ (AUC = 0.91, p=0.004). The resection ratio of spk-HFOs in the 90-minute data (the spk-HFO detector developed from 10-minute data was applied to 90-minute data) yielded an AUC of 0.88, which was comparable to the use of resection ratio of eHFOs.

**Figure 5.**
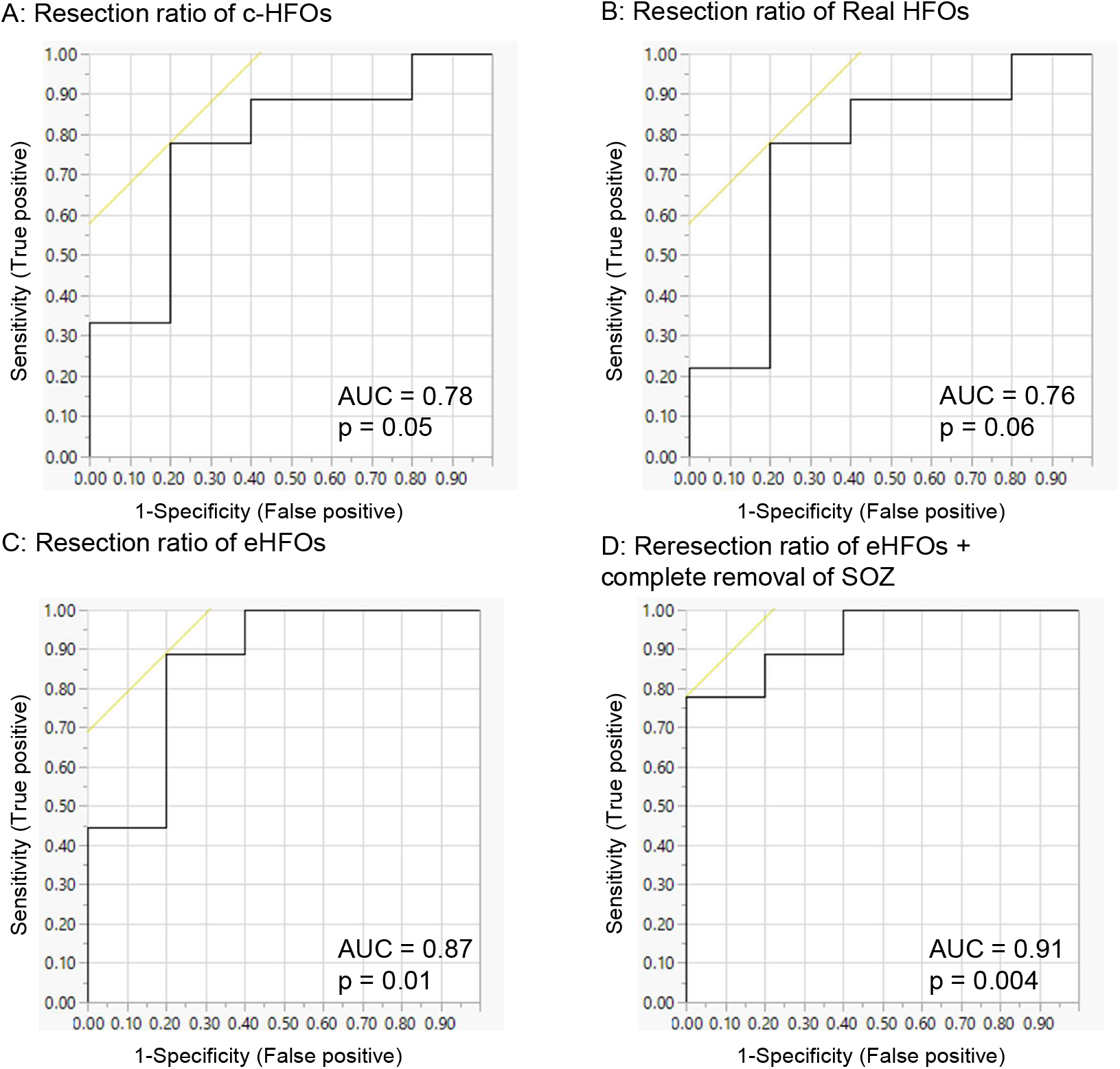
The accuracy of models incorporating HFO resection ratio. We constructed post-operative seizure outcome prediction models using HFO resection ratio derived from 90-minute EEG data (n = 14). Each receiver-operating characteristics (ROC) curve delineates the accuracy of seizure outcome classification of a given model, using the area under the ROC curve statistics. (A) HFO resection ratio using c-HFOs (raw HFO detections) was used as a single classifier. (B) Real HFO (rejection of artifacts from c-HFOs using the deep-learning based artifact detector) resection ratio was used as a single classifier. (C) eHFO (using the reverse engineering approach) resection ratio was used as a single classifier, which showed significant improvement in the prediction. (D) A multiple regression model incorporating the resection ratio of eHFOs and complete removal of the SOZ (yes or no) was used, which demonstrated further improved predictive value of post-operative seizure outcomes. HFO: High-frequency oscillation; c-HFOs: candidate HFOs; eHFOs: epileptogenic HFOs; SOZ: Seizure onset zone.

### Characterization of eHFOs and non-eHFOs using the time-frequency map and perturbation analysis

#### Time-frequency plot characteristics of epileptogenic and non-epileptogenic HFOs

The analysis of the time-frequency map demonstrated that eHFOs had higher values throughout the frequency band, including both ripples (80-250 Hz) and fast ripples (250-500 Hz) around the center point (0 ms, where HFOs were detected) than non-eHFOs. There were statistically higher values of eHFOs at the low-frequency band throughout the time window compared to non-eHFOs (**Figure 6A, B**). This pattern resembles an inverted T-shaped, limited to −45 ms to +45 ms on the time axis and 10 Hz to 59 Hz on the frequency axis. The mean pixel values (value = 1, if p-value < 0.005 from one-tailed t-test; value = 0, otherwise) were not different between ripples and fast ripples as a group (mean = 0.48 vs. 0.32; p=0.15, two-tailed t-test). In individual analysis, statistically significant pixel values were present in 12/15 subjects for ripples and 12/15 subjects for fast ripples (details in **Supplementally Table**).

**Figure 6.**
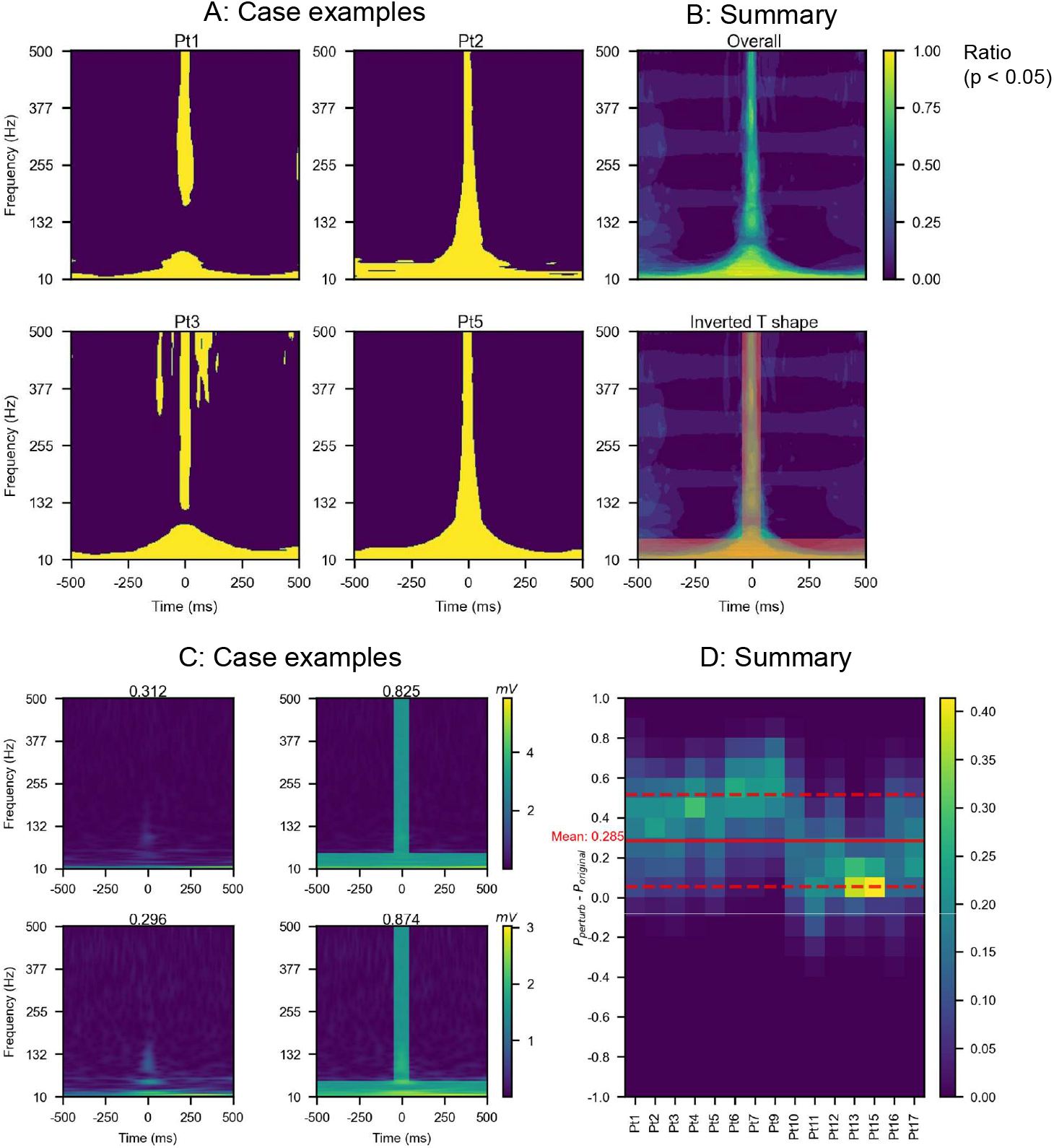
Characteristics in the time-frequency plot of eHFOs against non-eHFOs. (A) The time-frequency plot characteristics of epileptogenic and non-epileptogenic HFOs for Pt 1, 2, 3, and 5. The yellow-colored regions in the figure stood for the pixels, where the power spectrum of eHFOs is statistically higher than (p-value below 0.005 from one-tailed t-test) non-eHFOs. The figure showed one set of clearly interpretable distinguishing features between eHFOs and non-eHFOs: the eHFOs generally have higher power at higher frequencies during the HFO event (center part along the time axis), and more power in the low-frequency region in the entire time interval. Panel (B-top) was generated by taking the average of the individual binary images from each of the 15 patients. It showed the distinguishing features are also significant at the population level. The inverted T-shaped was designed to approximate the differentiating region on the time-frequency plot (B-bottom). (C) The model’s response to the inverted T-shaped perturbation on time-frequency plot. We provide two examples for perturbation for non-eHFO events in Pt 12. Each row presents one example and the first column indicates the original timefrequency plot while the second indicates the perturbed time-frequency plot based on the inverted T-shaped perturbation. The prediction value of the model changed from below 0.4 (therefore originally labeling it as non-eHFO) to above 0.8 (thus a change of +0.4) implying that the perturbed HFO would correspond to an eHFO. (D) The change in model confidence in population level. Each column (along the y-axis) is a histogram of the change in confidence for one distinct patient. It shows the frequency distribution of confidence changes after adding the inverted T-shaped perturbation to the time-frequency plot to all classified non-eHFOs for the given patient. The change in confidence level is significant, with an average of 0.285 noted as the red solid line in the histogram (a standard deviation noted as the red dashed line). HFO: High-frequency oscillation; eHFOs: epileptogenic HFOs; non-eHFOs: non-epileptogenic HFOs

#### Perturbation analysis to investigate salient features of epileptogenic and non-epileptogenic HFOs

By utilizing the inverted T-shaped template found in the time-frequency map, we observed that the inverted T-shaped perturbation on the time-frequency plot significantly increased the model prediction probability towards eHFOs (mean probability increase was 0.285, p < 0.001) (**Figure 6C, D**). Based on the positive correlation between eHFOs and spk-HFOs, we set our hypothesized TDCP pattern as a spike. Therefore, we analyzed the effect of introducing a spike-like shape in the amplitude-coding plot. By introducing a downgoing or an upgoing spike close to 0 ms location in a non-eHFOs event, the model confidence increased towards an eHFO event (**Figure 7A, D**). On the population level, the time step with the highest probability increase was around the center location along the time-axis (**Figure 7B, E**), where the HFO was detected. Moreover, the prevalent probability increase in non-eHFO events among all 15 patients who underwent resection (**Figure 7C, F**) demonstrated the non-trivial model response by introducing a spike in the time domain (mean probability increase of 0.438 for a downgoing spike introduction, and 0.465 for an upgoing spike introduction, both with p< 0.001).

**Figure 7.**
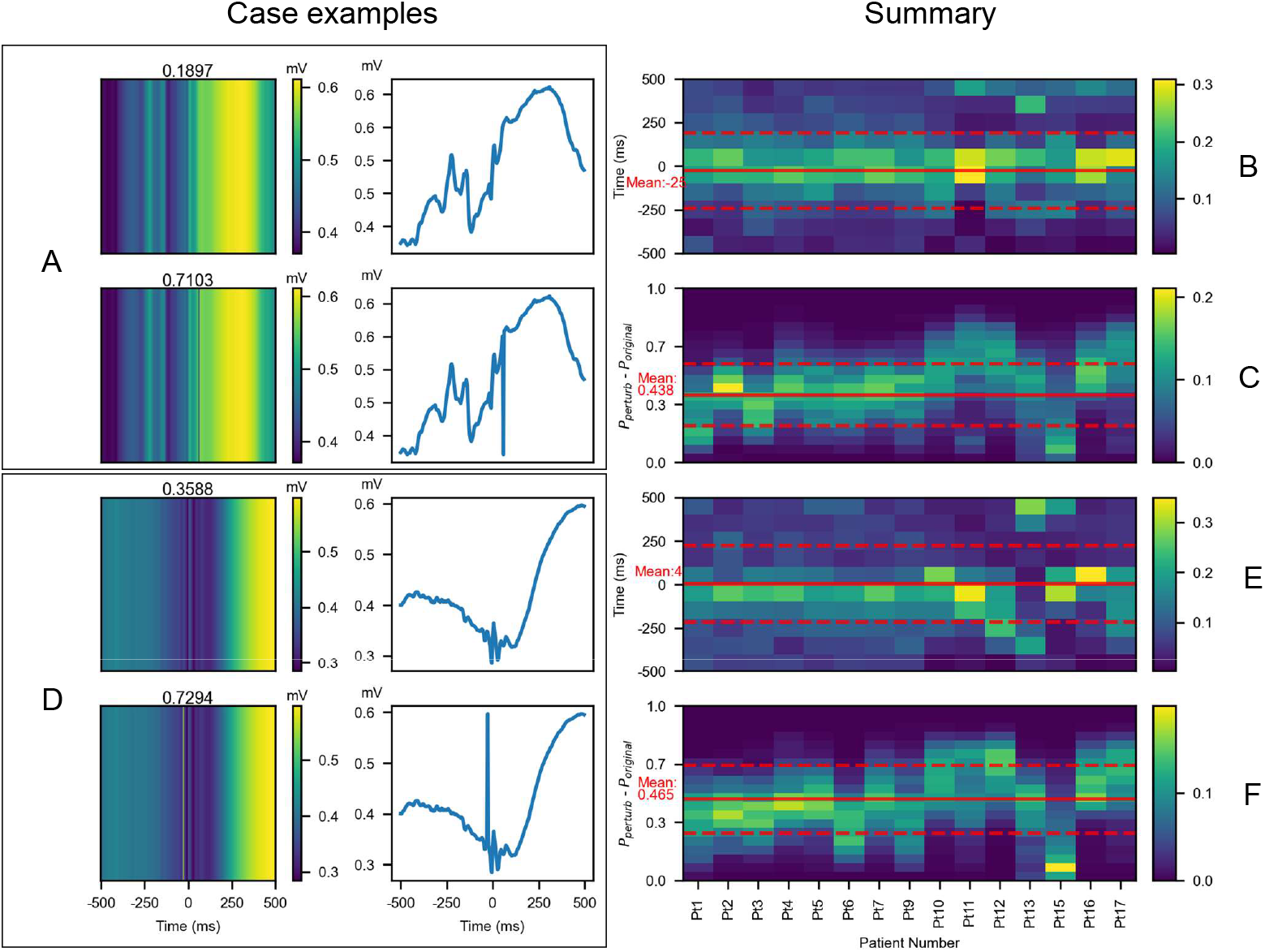
The model’s responses to injecting a spike-like feature into the amplitude-coding plot. (A,D) Examples of introducing a downgoing (A) and upgoing (D) spike feature in classified non-eHFO events. These demonstrate that on the introduction of a spike-like perturbation, the model predicts higher confidence toward eHFOs. Subfigure A shows the original amplitude encoding image input and the corresponding time-series signal (top row), and the perturbed amplitude-coding plot, and the corresponding time-series signal with downgoing spike perturbation (bottom row). Similarly, subfigure D shows the same information about a different classified eHFO but with upgoing spike perturbation. (B, C) For each non-eHFO, a downgoing spike perturbation could be introduced at every point in the time interval. For the perturbation, which resulted in the maximum change in confidence, its location (relative to the center, thus in the range of +500 to −500 ms) and the resulting change in confidence (−0.5, 1) are noted. For each patient, we compute a histogram for the distribution of the change in confidence (C) and a separate histogram for the location of the spike over non-eHFOs (B). The same steps are repeated for upgoing spike perturbation, and the results are shown in (E, F). For both up-and-downgoing spikes, the histograms (B, E) show that the spikes located close to the HFO event lead to maximum change in confidence. The change in confidence for both up-and-downgoing spike perturbation is significantly greater than zeros, with means downgoing: 0.438 and upgoing: 0.465, and the locations leading to the max perturbation are close to zeros location with means downgoing: −25ms and upgoing: +4ms. All of these means are noted as red solid lines in each histogram (a standard deviation noted as red dashed lines). HFO: High-frequency oscillation; eHFOs: epileptogenic HFOs; non-eHFOs: non-epileptogenic HFOs

## DISCUSSION

We demonstrated how DL might be used in HFO classification to complement experts at multiple levels. As a first step, we demonstrated that a DL-based algorithm robustly emulated HFO annotations by human experts in rejecting artifacts and determining if an event is associated with a spike. Thus, the associated DL models could help experts with their manual verifications and significantly reduce experts’ efforts without compromising performance. More importantly, in the second step, we showed how DL could create novel computational models which could define morphological classes of HFOs indicative of eHFOs and non-eHFOs when guided only by clinical outcomes such as seizure outcomes and resection status of the channels without any expert EEG labeling. We took advantage of our large dataset containing more than 30,000 HFOs within 90 minutes of EEG data from subjects with known post-operative seizure outcomes. The model was trained with a novel weakly supervised approach that utilized the inexact label of resected and non-resected status to discover eHFOs and non-eHFOs. We further showed that the DL-defined eHFOs were clinically relevant and possessed two salient features (HFOs associated with spikes and the inverted T-shaped pattern in scalogram) that experts had identified as characteristics of eHFOs. The removal rate of eHFOs correlated to seizure-freedom after resection, but no such relationship was seen with non-eHFOs. The use of the removal ratio of eHFOs seems to have an additional value on the status of complete removal of the SOZ, but the number of patients may be too small to conclude this definitively. By comparing the predicted results with expert annotation, we observed that the most classified eHFOs were spk-HFOs. Similarly, most non-eHFOs were non-spk-HFOs. At SOZ, we observed a higher rate of eHFOs compared to that of non-eHFOs.

We next elaborated on how the traits of DL-discovered eHFOs compared with hypothesized traits of so-called pathological HFOs. Pathological HFOs, often seen as fast ripples (FRs: 250 Hz or above), are believed to be excitatory neuronal activities, such as summated action potentials from synchronously bursting neurons.^6, 46^ Contrarily, physiological HFOs, which often involve ripples (80-250 Hz), are considered to reflect summated inhibitory postsynaptic potentials.^40, 47^ These biological mechanisms are hypothesized to lead to morphological differences between pathological and physiological HFOs, and hence, making them discoverable with computer vision. We used the trained DL model to derive certain signatures of eHFOs in the time-frequency domain that agreed with such common clinical knowledge. In particular, based on the analysis of time-frequency plots of eHFOs and non-eHFOs, eHFOs showed stronger signals throughout ripple and fast ripple bands at the onset of HFOs, which shares the similar observation in HFOs seen at the SOZ.^48^ Meanwhile, eHFOs generally showed stronger signals in the low frequency region throughout the time window, which might represent an inhibitory slow-wave postsynaptic component, coupled with out-of-phase excitatory fast firing of HFOs.^49^ Taking advantage of these observations, we designed perturbations in the input to probe whether these characteristics were actually the salient features that the model relied on to make a prediction. The group analysis on 90 min data demonstrated that both characteristics led to a significant model confidence increase towards eHFOs. Similarly, we found that the model had automatically learned the salient characteristics of manually labeled spk-HFOs. In addition, the artificial introduction of spike-like activity around the HFO onset also increased model confidence towards eHFOs. Clearly, such expert knowledge was not hard-coded into the DNNs and was automatically inferred only from the partial clinical outcomes. It is noteworthy that we discovered such salient features of epileptogenic HFOs from exploratory approaches utilizing DL. Clearly, one cannot exhaustively enumerate all the features that the DL model used to make its predictions, and these features are only some of the salient ones that aligned well with expert knowledge. As the study grows in scale and the DL models become accurate and robust in discovering eHFOs, we expect new salient characteristics to be discovered. The prior studies used DL architecture to classify HFOs using human-annotated data,^36, 37^ however, such approach has constraints including necessity of human experts and its associated potentially unfavorable inter-rater reliability among experts.^23^ This is the first study to demonstrate that DL algorithms can learn from clinical outcomes (such as seizure-freedom after resection) and soft labels (such as resected vs. preserved status). Our approach expands the spectrum of clinical usage, since there will be no need for expert annotation and easily applied to a larger dataset. In this study, we determined the resection status of channels among human experts, by reviewing pre-and post-resection intraoperative picture and post-operative brain MRI. This subjective approach might provide another type of labeling error, thus automated and objective determination of channel resection status might have been considered.

This work indicates that our deep learning approach can overcome issues including poor inter-rater reliability of HFO classification among human experts and their time constraints of analysis. Although our interrater reliability in this study was favorable as previously reported,^50^ we expect the agreement will likely diminish when raters are from different institutions or have different experience levels.^15, 23^ Using HFO analysis, we may identify brain tissue that needs to be removed during surgery without human experts’ effort if an algorithm was trained from large enough EEG data from patients with known post-operative outcomes. Notably, validation was performed using patient-wise cross-validation to maintain generalizability across patients so that the results may be applicable to new patients.

There are several limitations in our study. Regarding the dataset, we have included only 19 patients, and there were only 14 patients who had resective surgery with known clinical outcomes at 24 months. Although we analyzed extended EEG data (90 minutes) from nine patients who achieved seizure-freedom at 24 months after surgery, we did not analyze the entire EEG recordings. It is of interest to include more patients and EEG data to train our DL algorithm and evaluate how much the performance could improve. In fact, we are planning to build a model using EEG data from more than 100 patients who underwent resection. Also, all the data were from pediatric patients from the same institution. With a diversified age range and epilepsy pathology, the morphology of HFOs may change, and the algorithm might be trained differently.

In this work, we proposed an automated tool to analyze HFOs in a large dataset by using DL to simulate a human expert’s visual verification and to predict seizure-free status through an outcome-based reverse engineering approach. Future work to further refine this methodology by examining more datasets from multiple institutions is still needed. By refining a method to distinguish epileptogenic HFOs from others, we will be better positioned to confidently use HFOs to guide resection margin in clinical trials to improve the chance of post-operative seizure-freedom in patients with drug-resistant epilepsy who undergo epilepsy surgery.

## ACKNOWLEDGMENT

The authors have no conflict of interest to disclose. HN is supported by the Susan Spencer Clinical Research Training Fellowship in Epilepsy from the American Academy of Neurology (#20184605), the Pediatric Victory Foundation, the Sudha Neelakantan & Venky Harinarayan Charitable Fund, the Elsie and Isaac Fogelman Endowment, and the UCLA Children’s Discovery and Innovation Institute (CDI) Junior Faculty Career Development Grant (#CDI-SEED-010121). SAH has received research support from the Epilepsy Therapy Project, the Milken Family Foundation, the Hughes Family Foundation, the Elsie and Isaac Fogelman Endowment, Eisai, Lundbeck, Insys, Zogenix, GW Pharmaceuticals, UCB, and has received honoraria for service on the scientific advisory boards of Questcor, Mallinckrodt, Insys, UCB, and Upsher-Smith, for service as a consultant to Eisai, UCB, GW Pharmaceuticals, Insys, and Mallinckrodt, and for service on the speakers’ bureaus of Mallinckrodt and Greenwich Bioscience. RS serves on scientific advisory boards and speakers bureaus for and has received honoraria and funding for travel from Eisai, Greenwich Biosciences, UCB Pharma, Sunovion, Supernus, Lundbeck Pharma, Liva Nova, and West Therapeutics (advisory only); receives royalties from the publication of Pellock’s Pediatric Neurology (Demos Publishing, 2016) and Epilepsy: Mechanisms, Models, and Translational Perspectives (CRC Press, 2011). JEJ is supported by the National Institute of Neurological Disorders and Stroke (NINDS) U54NS100064 and R01NS033310. The research described was also supported by NIH/National Center for Advancing Translational Science (NCATS) UCLA CTSI Grant Number UL1TR001881.

We are indebted to Jason T. Lerner, Joyce H. Matsumoto, Lekha M. Rao, Rajsekar R. Rajaraman, Jimmey C Nguyen, Maria Garcia Roca, Richard Le, Patrick Wilson, and Kristina Murata for their assistance in the study and sample acquisition. We also greatly thank Ms. Shelly Recchio for her assistant in the graphical abstract.

## Supplementary Figure

**Supplementary Figure.**
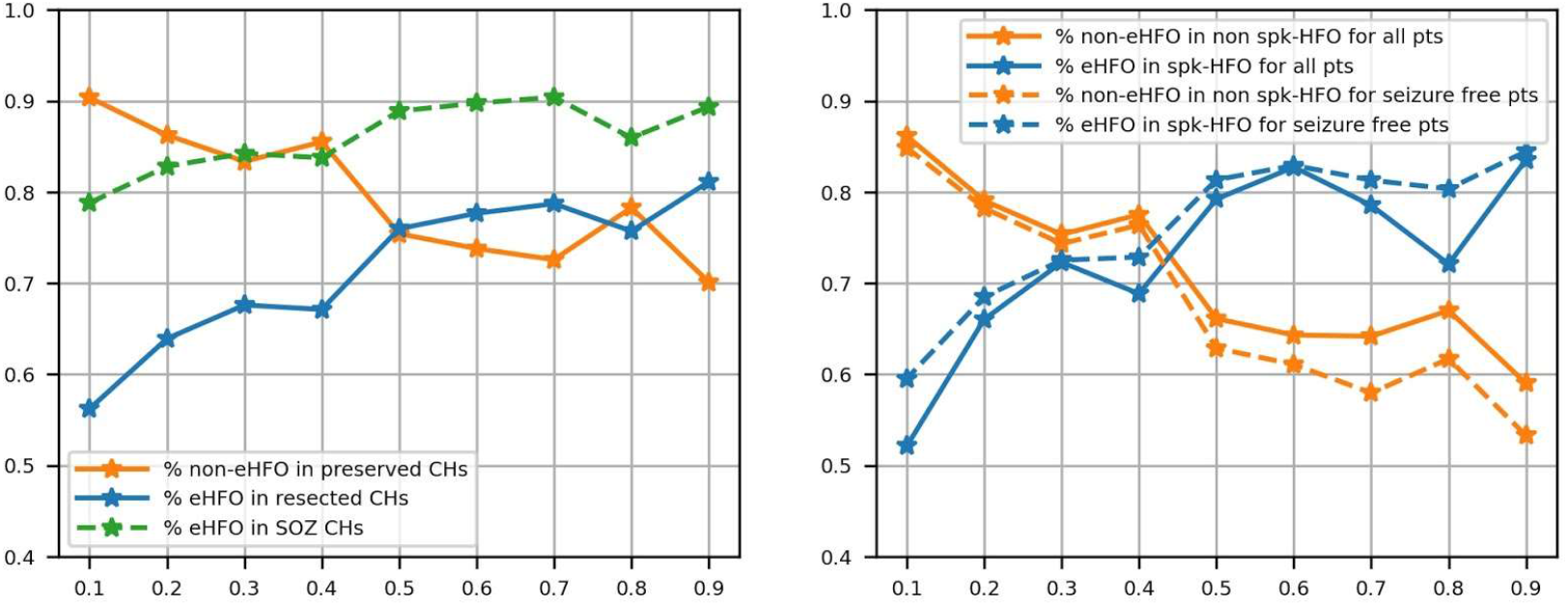
To find the optimal weight (w) term in the Binary Cross Entropy Loss, we swept the w from 0.1 to 0.9 and compared the performance of the model trained by each w with our hypothesis. The first criterion for selecting an optimal weight w was based on the following hypothesis that requires no expert input: For post-surgery seizure-free patients, the majority of the classified non-eHFOs should be located within the preserved regions, and correspondingly, the majority of the eHFOs should be located within resected regions; moreover, a vast majority of the HFOs in the SOZ channels should be labeled as eHFOs. Thus, an optimal w should balance the first two percentages (i.e., % of non-eHFOs in preserved regions and the % of eHFOs in resected regions) and maximize the percentage of eHFOs in the SOZ channel. As the Left plot shows, w =0.5 satisfies this criterion: The % of eHFOs in the resected CHs (orange) equals the % of non-eHFOs in preserved channels (blue), and the % of eHFOs in SOZ CHs reaches a maximum value. A second criterion for validating an optimal choice of w was based on the hypothesis that there should be an agreement between the data-driven predictions and expert knowledge: For all patients, the majority of the detected eHFOs should also be labeled by experts as pathological HFOs (i.e., spk-HFOs), and correspondingly, the majority of the detected non-eHFOs should be labeled as physiological by the experts (i.e., non-spk-HFOs). An optimal w would be at a value where the corresponding percentages are balanced. The Right figure shows that the respective percentages (orange vs. blue) become equal (~72%) at w= 0.45 and remain still balanced at w=0.5 (~78% vs. ~68). Given potential inter-rater variations that could change the percentages, we considered w=0.5 to be an optimal choice for our study. HFO: High-frequency oscillation; eHFOs: epileptogenic HFOs; non-eHFOs: non-epileptogenic HFOs; SOZ: Seizure onset zone

---

**Supplementary Table.** Mean pixel statistical value at the HFO event onset (−45ms to +45ms)

| Pt. No. | Ripples | Fast ripples |  |
| --- | --- | --- | --- |
| 1 | 0.35 | 0.57 |  |
| 2 | 0.82 | 0.41 |  |
| 3 | 0.41 | 0.50 |  |
| 4 | 0.00 | 0.00 |  |
| 5 | 0.85 | 0.50 |  |
| 6 | 0.76 | 0.28 |  |
| 7 | 0.33 | 0.03 |  |
| 9 | 0.00 | 0.17 |  |
| 10 | 0.68 | 0.67 |  |
| 11 | 0.04 | 0.00 |  |
| 12 | 0.71 | 0.39 |  |
| 13 | 0.00 | 0.00 |  |
| 15 | 0.77 | 0.22 |  |
| 16 | 0.96 | 0.86 |  |
| 17 | 0.50 | 0.26 |  |
| Overall | 0.48 | 0.32 | *p = 0.15 |

\*Two-tailed t-test

## Notes

### Competing Interest Statement

The authors have declared no competing interest.

## REFERENCES

1. Kwan P, Brodie MJ. Early identification of refractory epilepsy. The New England journal of medicine 2000;342:314–319.

2. Wiebe S, Blume WT, Girvin JP, Eliasziw M. A randomized, controlled trial of surgery for temporal-lobe epilepsy. The New England journal of medicine 2001;345:311–318.

3. Engel J, Jr., McDermott MP, Wiebe S, et al. Early surgical therapy for drug-resistant temporal lobe epilepsy: a randomized trial. JAMA 2012;307:922–930.

4. Asano E, Juhasz C, Shah A, Sood S, Chugani HT. Role of subdural electrocorticography in prediction of long-term seizure outcome in epilepsy surgery. Brain 2009;132:1038–1047.

5. Dwivedi R, Ramanujam B, Chandra PS, et al. Surgery for Drug-Resistant Epilepsy in Children. The New England journal of medicine 2017;377:1639–1647.

6. Bragin A, Engel J, Jr., Wilson CL, Fried I, Mathern GW. Hippocampal and entorhinal cortex high-frequency oscillations (100--500 Hz) in human epileptic brain and in kainic acid--treated rats with chronic seizures. Epilepsia 1999;40:127–137.

7. Bragin A, Engel J, Jr., Wilson CL, Vizentin E, Mathern GW. Electrophysiologic analysis of a chronic seizure model after unilateral hippocampal KA injection. Epilepsia 1999;40:1210–1221.

8. Staba RJ, Wilson CL, Bragin A, Jhung D, Fried I, Engel J, Jr. High-frequency oscillations recorded in human medial temporal lobe during sleep. Annals of neurology 2004;56:108–115.

9. Jirsch JD, Urrestarazu E, LeVan P, Olivier A, Dubeau F, Gotman J. High-frequency oscillations during human focal seizures. Brain 2006;129:1593–1608.

10. Worrell GA, Parish L, Cranstoun SD, Jonas R, Baltuch G, Litt B. High-frequency oscillations and seizure generation in neocortical epilepsy. Brain 2004;127:1496–1506.

11. Jacobs J, Zijlmans M, Zelmann R, et al. High-frequency electroencephalographic oscillations correlate with outcome of epilepsy surgery. Annals of neurology 2010;67:209–220.

12. Wu JY, Sankar R, Lerner JT, Matsumoto JH, Vinters HV, Mathern GW. Removing interictal fast ripples on electrocorticography linked with seizure freedom in children. Neurology 2010;75:1686–1694.

13. Akiyama T, McCoy B, Go CY, et al. Focal resection of fast ripples on extraoperative intracranial EEG improves seizure outcome in pediatric epilepsy. Epilepsia 2011;52:1802–1811.

14. van’t Klooster MA, van Klink NEC, Zweiphenning W, et al. Tailoring epilepsy surgery with fast ripples in the intraoperative electrocorticogram. Annals of neurology 2017;81:664–676.

15. Jacobs J, Wu JY, Perucca P, et al. Removing high-frequency oscillations: A prospective multicenter study on seizure outcome. Neurology 2018;91:e1040–e1052.

16. Matsumoto A, Brinkmann BH, Matthew Stead S, et al. Pathological and physiological high-frequency oscillations in focal human epilepsy. J Neurophysiol 2013;110:1958–1964.

17. Melani F, Zelmann R, Mari F, Gotman J. Continuous High Frequency Activity: a peculiar SEEG pattern related to specific brain regions. Clinical neurophysiology: official journal of the International Federation of Clinical Neurophysiology 2013;124:1507–1516.

18. Frauscher B, von Ellenrieder N, Zelmann R, et al. High-Frequency Oscillations in the Normal Human Brain. Annals of neurology 2018;84:374–385.

19. Benar CG, Chauviere L, Bartolomei F, Wendling F. Pitfalls of high-pass filtering for detecting epileptic oscillations: a technical note on “false” ripples. Clinical neurophysiology: official journal of the International Federation of Clinical Neurophysiology 2010;121:301–310.

20. Nonoda Y, Miyakoshi M, Ojeda A, et al. Interictal high-frequency oscillations generated by seizure onset and eloquent areas may be differentially coupled with different slow waves. Clinical neurophysiology: official journal of the International Federation of Clinical Neurophysiology 2016;127:2489–2499.

21. Weiss SA, Orosz I, Salamon N, et al. Ripples on spikes show increased phase-amplitude coupling in mesial temporal lobe epilepsy seizure-onset zones. Epilepsia 2016;57:1916–1930.

22. Motoi H, Miyakoshi M, Abel TJ, et al. Phase-amplitude coupling between interictal high-frequency activity and slow waves in epilepsy surgery. Epilepsia 2018;59:1954–1965.

23. Spring AM, Pittman DJ, Aghakhani Y, et al. Interrater reliability of visually evaluated high frequency oscillations. Clinical neurophysiology: official journal of the International Federation of Clinical Neurophysiology 2017;128:433–441.

24. Cimbalnik J, Brinkmann B, Kremen V, et al. Physiological and pathological high frequency oscillations in focal epilepsy. Ann Clin Transl Neurol 2018;5:1062–1076.

25. Nevalainen P, von Ellenrieder N, Klimeš P, Dubeau F, Frauscher B, Gotman J. Association of fast ripples on intracranial EEG and outcomes after epilepsy surgery. Neurology 2020;95:e2235–e2245.

26. Nariai H, Hussain SA, Bernardo D, et al. Prospective observational study: Fast ripple localization delineates the epileptogenic zone. Clinical neurophysiology: official journal of the International Federation of Clinical Neurophysiology 2019;130:2144–2152.

27. Motoi H, Jeong JW, Juhasz C, et al. Quantitative analysis of intracranial electrocorticography signals using the concept of statistical parametric mapping. Sci Rep 2019;9:17385.

28. Kuroda N, Sonoda M, Miyakoshi M, et al. Objective interictal electrophysiology biomarkers optimize prediction of epilepsy surgery outcome. Brain Commun 2021;3:fcab042.

29. Jrad N, Kachenoura A, Merlet I, Nica A, Benar CG, Wendling F. Classification of high frequency oscillations in epileptic intracerebral EEG. Annu Int Conf IEEE Eng Med Biol Soc 2015;2015:574–577.

30. Amiri M, Lina JM, Pizzo F, Gotman J. High Frequency Oscillations and spikes: Separating real HFOs from false oscillations. Clinical neurophysiology: official journal of the International Federation of Clinical Neurophysiology 2016;127:187–196.

31. Jrad N, Kachenoura A, Merlet I, et al. Automatic Detection and Classification of High-Frequency Oscillations in Depth-EEG Signals. IEEE Trans Biomed Eng 2017;64:2230–2240.

32. Chaibi S, Lajnef T, Samet M, Jerbi K, Kachouri A. Detection of High Frequency Oscillations (HFOs) in the 80-500 Hz range in epilepsy recordings using decision tree analysis. International Image Processing, Applications and Systems Conference 2014:1–6.

33. Liu S, Gurses C, Sha Z, et al. Stereotyped high-frequency oscillations discriminate seizure onset zones and critical functional cortex in focal epilepsy. Brain 2018;141:713–730.

34. Lundervold AS, Lundervold A. An overview of deep learning in medical imaging focusing on MRI. Z Med Phys 2019;29:102–127.

35. Jing J, Sun H, Kim JA, et al. Development of Expert-Level Automated Detection of Epileptiform Discharges During Electroencephalogram Interpretation. JAMA neurology 2020;77:103–108.

36. Zhao B, Hu W, Zhang C, et al. Integrated Automatic Detection, Classification and Imaging of High Frequency Oscillations With Stereoelectroencephalography. Front Neurosci 2020;14:546.

37. Zuo R, Wei J, Li X, et al. Automated Detection of High-Frequency Oscillations in Epilepsy Based on a Convolutional Neural Network. Front Comput Neurosci 2019;13:6.

38. Karimi D, Dou H, Warfield SK, Gholipour A. Deep learning with noisy labels: Exploring techniques and remedies in medical image analysis. Med Image Anal 2020;65:101759.

39. Nakai Y, Jeong JW, Brown EC, et al. Three- and four-dimensional mapping of speech and language in patients with epilepsy. Brain 2017;140:1351–1370.

40. Staba RJ, Wilson CL, Bragin A, Fried I, Engel J, Jr. Quantitative analysis of high-frequency oscillations (80-500 Hz) recorded in human epileptic hippocampus and entorhinal cortex. J Neurophysiol 2002;88:1743–1752.

41. Navarrete M, Alvarado-Rojas C, Le Van Quyen M, Valderrama M. RIPPLELAB: A Comprehensive Application for the Detection, Analysis and Classification of High Frequency Oscillations in Electroencephalographic Signals. PLoS One 2016;11:e0158276.

42. He K, Zhang X, Ren S, Sun J. Deep residual learning for image recognition. Proceedings of the IEEE conference on computer vision and pattern recognition 2016:770–778.

43. Tamilia E, Park EH, Percivati S, et al. Surgical resection of ripple onset predicts outcome in pediatric epilepsy. Annals of neurology 2018;84:331–346.

44. Guidotti R, Monreale A, Ruggieri S, Turini F, Giannotti P, Pedreschi D. A survey of methods for explaining black box models. ACM computing surveys (CSUR) 2018;51:1–42.

45. Selvaraju R, Cogswell M, Das A, Vedantam R, Parikh D, Batra D. Grad-CAM: Visual explanations from deep networks via gradient-based localization. Proceedings of the IEEE international conference on computer vision 2017:618–626.

46. Bragin A, Mody I, Wilson CL, Engel J, Jr. Local generation of fast ripples in epileptic brain. The Journal of neuroscience: the official journal of the Society for Neuroscience 2002;22:2012–2021.

47. Buzsáki G, Horváth Z, Urioste R, Hetke J, Wise K. High-frequency network oscillation in the hippocampus. Science 1992;256:1025–1027.

48. Charupanit K, Sen-Gupta I, Lin JJ, Lopour BA. Amplitude of high frequency oscillations as a biomarker of the seizure onset zone. Clinical neurophysiology: official journal of the International Federation of Clinical Neurophysiology 2020;131:2542–2550.

49. Jefferys JG, Menendez de la Prida L, Wendling F, et al. Mechanisms of physiological and epileptic HFO generation. Prog Neurobiol 2012;98:250–264.

50. Nariai H, Wu JY, Bernardo D, Fallah A, Sankar R, Hussain SA. Interrater reliability in visual identification of interictal high-frequency oscillations on electrocorticography and scalp EEG. Epilepsia open 2018;3:127–132.

